# Threat induces changes in cardiac activity and metabolism negatively impacting survival in flies

**DOI:** 10.1101/2020.12.02.408161

**Authors:** Natalia Barrios, Matheus Farias, Marta A Moita

## Abstract

Adjusting to a dynamic environment involves fast changes in the body’s internal state, characterized by coordinated alterations in brain activity, physiological and motor responses. Threat-induced defensive states are a classic example of coordinated adjustment of bodily responses, being cardiac regulation one of the best characterized in vertebrates. A great deal is known regarding the neural basis of invertebrate defensive behaviours, mainly in *Drosophila melanogaster*. However, whether physiological changes accompany these remains unknown. Here, we set out to describe the internal bodily state of fruit flies upon an inescapable threat and found cardiac acceleration during running and deceleration during freezing. In addition, we found that freezing leads to increased cardiac pumping from the abdomen towards the head-thorax, suggesting mobilization of energy resources. Concordantly, threat-triggered freezing reduces sugar levels in the hemolymph and renders flies less resistant to starvation. The cardiac responses observed during freezing were absent during spontaneous immobility, underscoring the active nature of freezing response. Finally, we show that baseline cardiac activity predicts the amount of freezing upon threat. This work reveals a remarkable similarity with the cardiac responses of vertebrates, suggesting an evolutionarily convergent defensive state in flies. Our findings are at odds with the widespread view that cardiac deceleration while freezing has first evolved in vertebrates and that it is energy sparing. Investigating the physiological changes coupled to defensive behaviours in the fruit fly has revealed that freezing is costly, yet accompanied by cardiac deceleration, and points to heart activity as a key modulator of defensive behaviours.

## Introduction

Internal states that accompany behaviours, emotions and cognitive processes include all bodily adjustments that facilitate short-term deviations from homeostasis thereby enabling efficient coping with significant challenges. Cardiac activity is widely used to evaluate the body’s internal state. A classic example of cardiac responses to environmental challenge is the tight coupling between heart rate and defensive behaviours [1–3] observed upon threat. In vertebrates, including humans, escapable or imminent threats typically induce flight-or-fight responses associated with increased heart rate, tachycardia [4–6]. By contrast, more distant or inescapable threats lead to freezing responses with decreased heart rate, bradycardia [1,7,8]. The above changes in cardiac activity during defensive behaviours are thought to be linked to metabolic needs of threatened animals, as the circulatory system regulates oxygen distribution throughout the body [4]. In addition to these sustained changes in cardiac activity, threat induces transient cardiac arrest accompanying orienting responses as part of the startle reflex [9]. The tight regulation of cardiac function upon threat is mediated by the autonomic nervous system that first evolved in vertebrates [10]. Whether life threatening situations, such as an encounter with a predator, leads to changes in cardiac activity and other physiological processes in invertebrates, which lack an autonomic nervous system, remains largely unexplored.

Insects show a large repertoire of defensive behaviours, including freezing and fleeing responses, upon detection of a threat [11–14]. The fruit fly has become a very useful model to study the neuronal circuits of defensive behaviours, from the integration of cues of threat and surrounding context, to the selection and execution of the defensive responses [11,14–16]. It has been previously postulated that threat induces a sustained change in the flies’ internal state, akin to a primitive emotional state [17]. However, the physiological changes that comprise this altered state have not been described. Here we aimed to test whether in the fruit fly these defensive behaviours are paralleled by physiological changes, focusing on cardiac activity during freezing and fleeing behaviours as well as during spontaneous locomotion and immobility.

Drosophila have an open circulatory system with a dorsal vessel, that extends from the posterior end of the abdomen to the head. The abdominal part of the dorsal vessel functions as a tubular contractile heart that periodically reverses the direction of beating. These cardiac reversals allow alternating between pumping the hemolymph towards the head-thorax or towards the abdomen [18]. As the circulation of blood in vertebrates, the flow of the hemolymph serves to transport immune cells, nutrients, and other metabolites between the thoracic and abdominal cavities. In contrast to vertebrates, where a crucial function of the circulatory system is the transport of oxygen to all parts of the body, insects possess a vast tracheal system covering the whole body that is responsible for oxygen transport. Yet, it has been demonstrated in the blowfly and other insects that cardiac reversals produce periodic pressure changes in thorax and abdomen resulting in expansion and contraction of tracheae and air sacs that support ventilation [19–21]. The small size of the fruit fly does not facilitate such a study, however a similar connection between the circulatory and respiratory systems has been hypothesized [22].

In this work, we develop a new technique to image adult cardiac activity in semi-restrained behaving flies. We found an astounding coupling between defensive behaviours and cardiac activity analogous to that found in vertebrates, i.e. running with heart acceleration of heart activity and freezing with heart deceleration. In addition, we found that freezing flies biased cardiac pumping towards the head and thorax, suggesting mobilization of resources from the abdomen. Concordantly, after sustained freezing flies show a decrease in sugar levels and they are less resistant to starvation. Finally, we observed that animals that will respond to threat by freezing or by fleeing have different patterns of cardiac reversal even before threat presentation. Using a regression model with cardiac reversal rate and cardiac reversal variability as predictors, we found that both variables are predictive of freezing intensity, establishing a link between cardiac function and the selection of defensive behaviours upon threat.

## Results

### Semi-restrained flies freeze or flee to repeated looming

Like vertebrates, unrestrained flies flexibly respond to an inescapable threat with freezing or flight attempts [11,13,14], allowing the study of the coupling of these behaviours with physiological adaptations. To assess whether cardiac activity is modulated by threat in a behaviour specific manner, we required flies to be tethered by the abdomen to a glass slide. We first tested whether these semi-restrained flies freeze and flee in response to looming stimuli, which mimic an object on collision course and reliably trigger defensive behaviours in a wide range of species including fruit flies [11,23,24]. Flies were dorsally tethered and placed on a ball while a computer monitor showed 20 repetitions of a looming black disc or a control visual stimulus, consisting of black dots randomly appearing on the blue screen over the course of a 5 min period (see Materials and Methods; Fig. 1 A, B). The behaviour of flies was recorded through an IR camera placed laterally. We found that semi-restrained flies freeze and flee upon looming (Movie S1 and S2). Jumps were also observed but not quantified in this work. To estimate the fly’s walking speed, we analysed the rotational movement of the ball using a custom written software in Bonsai [25].

**Figure 1.**
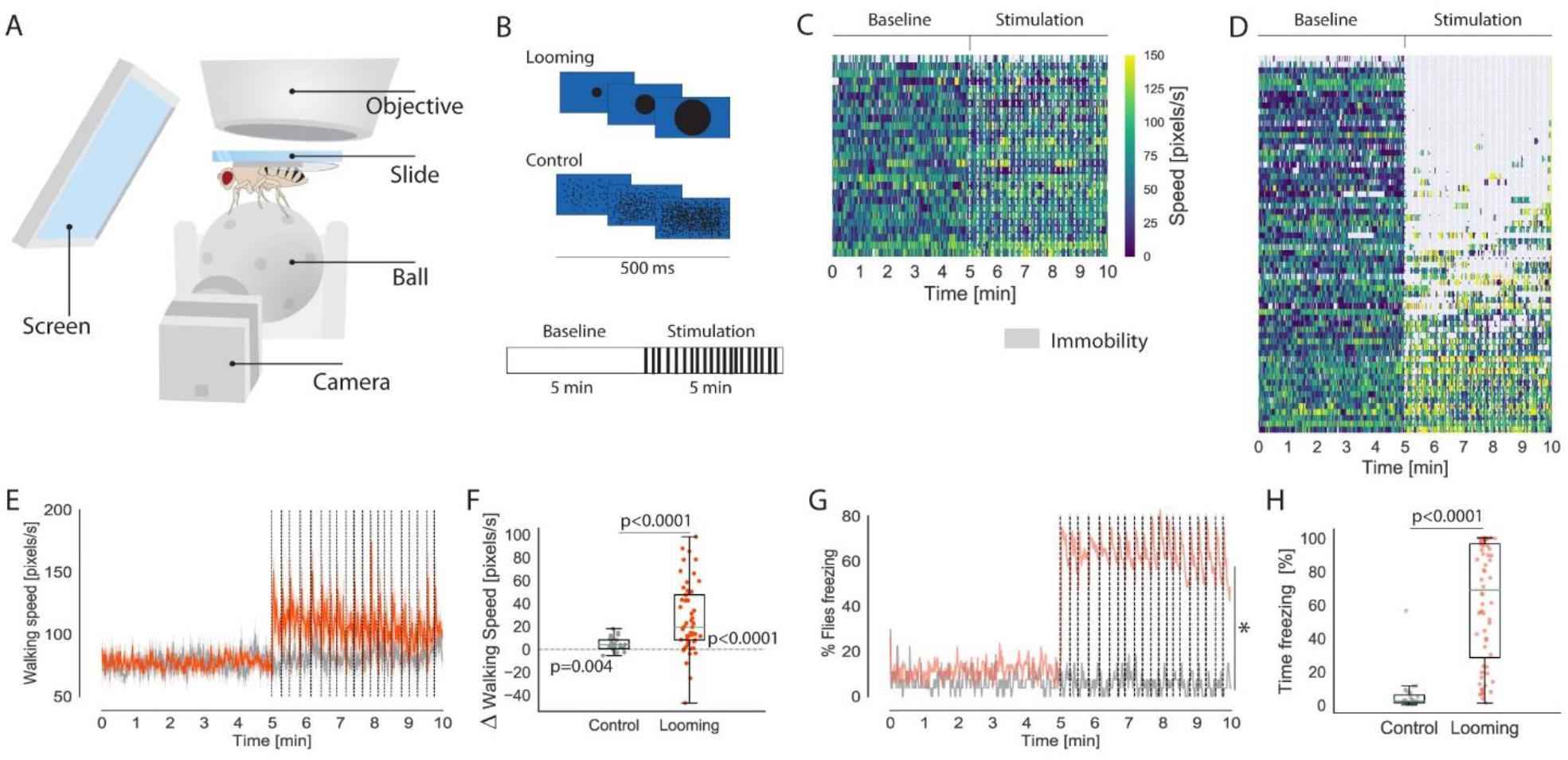
Semi-restrained flies flee or freeze upon looming presentation. (A) Schematic representation of the behavioural setup. (B) Schematic drawing of the visual stimuli and the protocol. After a 5-minute baseline flies were exposed to twenty 500 ms stimuli, every 10-20s, indicated by vertical black lines. (C-D) Walking speed (colour scale on the right) and freezing/immobility (grey) in 1 sec bins for control (C, n=17) and looming stimulated flies (D, n=64). Each row corresponds to one fly, rank ordered by maximum average walking speed during stimulation period. (E) Average walking speed (± s.e.m.) over the course of the experiment. (F) Change in walking speed per individual fly (each data point shows average speed during stimulation – average speed during baseline period). (G) Fraction of flies freezing over the course of the experiment. (H) Time spent freezing where each data point shows percent time freezing per individual fly. In E to H, grey correspond to control flies and red and pink to looming stimulated flies. Dashed vertical lines in C, D, E and G represent stimulus presentations. E-F include walking bouts only. In F and H and all figures hereafter, green horizontal lines represent medians, box represents interquartile range and whiskers min and max values.

We analysed locomotor behaviour excluding all immobility and grooming bouts, hence including only periods classified as walking (speed >30 pixels/s). The walking speed of flies increased during exposure to looming stimuli relative to baseline, during which walking speed remained constant (Wilcoxon test, p<0.0001; Fig. 1 C to F). Control stimulated flies also showed constant walking speed during baseline and an increase in walking speed during stimulation (Wilcoxon test, p=0.004; Figure 1 E, F) albeit lower than that observed in looming stimulated flies *(Mann-Whitney Utest,* p<0.0001).

To identify periods of immobility, we quantified pixel change in a region of interest surrounding the fly (see Methods, Fig. S1 A). Before stimulation, we found that flies were rarely immobile; when they were not walking they mostly groomed. In contrast, loomed animals, but not control ones, sustained long periods of immobility during the stimulation period (Fig. 1 C, D, G and H). One second after the last stimulus presentation, 75% of loomed flies (48/64) were freezing compared to 4%(1/27) of control stimulated flies *X*^2^ test, *p*< 0.0001; Fig. 1 G). It has been shown in enclosed arenas that the fraction of flies freezing increases gradually with each looming presentation [11]. Nevertheless, in our experimental conditions, the fraction of flies immobile stayed constant during the stimulation period; 75% (48/64) of flies were freezing one second after first loom and 68% (44/64) one second after last loom *(Χ^2^* test, p=0.677; Fig. 1 G). When we analysed the total amount of time each animal spent freezing during stimulation, we found that loomed animals froze more compared to control stimulated flies *(Mann-Whitney U test,* p<0.0001; Fig. 1 H). As shown for flies in enclosed arenas [11], we observed a bimodal distribution of the amount of time flies spend freezing during the period of exposure to looming stimuli (Fig. S1 B).

These data show that semi-restrained flies display both fleeing and freezing defensive responses upon threat, just as freely behaving flies in enclosed arenas do, allowing the search for possible coupling of cardiac function and defensive behaviours in the adult fruit fly.

### Flies show cardiac startle upon looming

The fruit fly’s heart is composed of two rows of cardiomyocytes that contract rhythmically, pumping the hemolymph alternatively in opposite directions: backward, towards the abdomen, or forward, towards the head and thorax [18] (Fig. 2 A). Alternation between backward and forward pumping is known as cardiac reversal.

**Figure 2.**
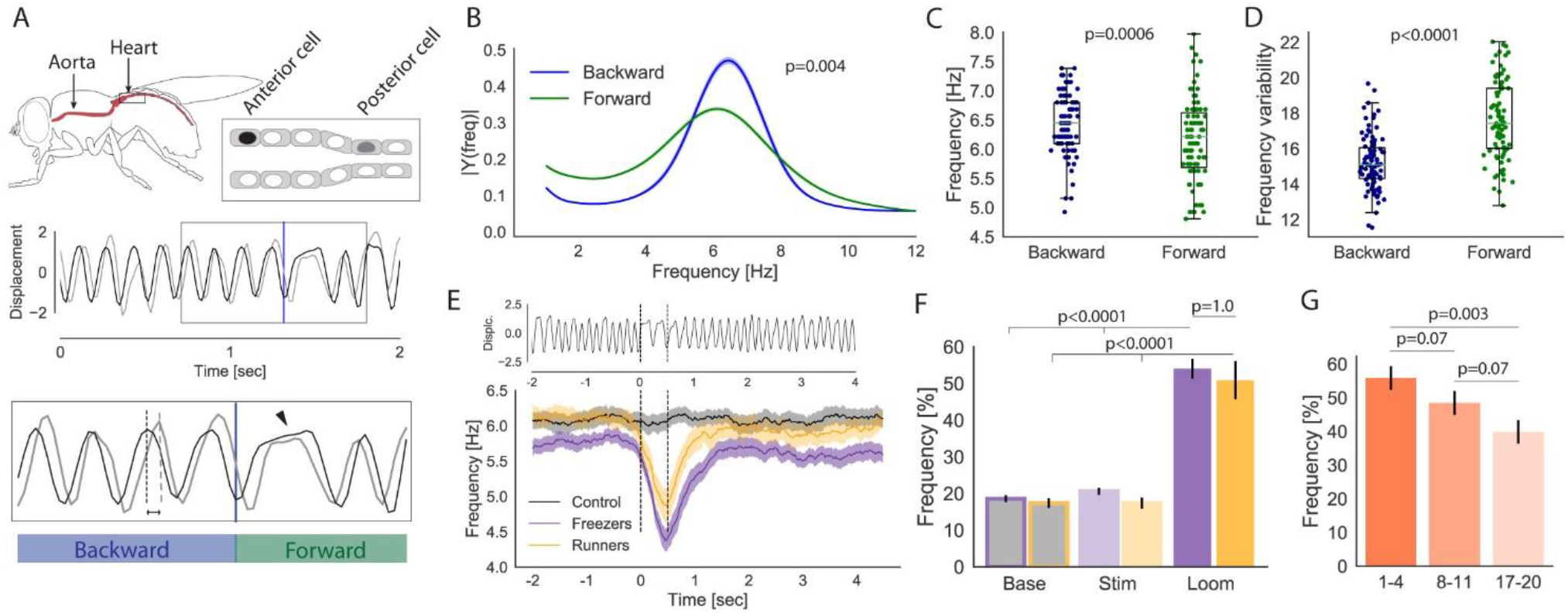
Looming triggers transient cardiac arrest in semi-restrained intact flies. (A) Illustration showing the dorsal vessel of the fruit fly (top-left) and a schematic representation of the cardiomyocytes of the contractile heart (top-right). Representative traces obtained by tracking the displacement of two cardiomyocytes, one more anterior and another more posterior (black and grey respectively) (middle row). Directional change of heartbeat analysed by metachrony of contractions (bottom row). The point arrow indicates a transient cardiac arrest associated with cardiac reversal. (B) Average power spectral density of backward and forward beating rates (± s.e.m.). (C-D) Frequency and variability of backward and forward beating rate per individual fly during the baseline period. (E) Representative cardiac trace showing a transient cardiac arrest during a looming stimulus (top). Average peak beating frequency (± s.e.m.), aligned to visual stimulus (bottom) for control flies and for runners and freezers. Dashed lines represent the beginning and end of the stimulus. (F) Frequency of cardiac arrest at random time points during baseline period, during stimulation period outside visual stimulus presentations, and during looming stimuli for freezer and runner flies (± s.e.m.). (G) Progression of cardiac arrest over the course of loom presentations. In E-F and hereafter, purple and orange correspond to freezers and runners respectively. In A-D and hereafter, blue and green correspond to backward and forward pumping respectively.

Cardiac activity has been analysed mainly in semi-intact preparations or anesthetized flies [26] and only a few studies have investigated its function in intact animals [22,27]. However, so far there are no studies showing the activity of the heart in behaving fruit flies. Here, we imaged the nuclei of cardiomyocytes expressing nls-dsRed in dorsally tethered awake flies placed on a ball (such that the head and legs move freely, Movie S3). To analyse cardiac activity we tracked the displacement, perpendicular to the longitudinal axis of the heart tube, of the nucleus of two cardiomyocytes, one in the second abdominal segment and another in the third one (Fig. 2 A). This allowed measuring both cardiac beating rate, using the displacement of one cardiomyocyte, and cardiac pumping direction, using the relative displacement of the two cardiomyocytes (see methods).

We first analysed the cardiac beating frequency during the baseline period. We performed Short-time Fast Fourier (SFF) on the signal from the anterior cell and found the peak beating frequency to be 6.33 ± 0.37 Hz (mode ± s.e.m.) which agrees with Klassen et al., 2017 [27]. It has been reported that the heart beats at different frequencies during backward and forward pumping bouts [22,28]. Thus, we analysed beating rate during backward and forward pumping separately. To this end, we concatenated all baseline backward bouts and all forward bouts for each animal and performed SFF on each concatenated signal. As reported previously [22], we found that during the baseline period backward beating is faster (Wilcoxon test, p=0.0006) and more regular than forward beating (Wilcoxon test, p<0.0001; Fig. 2 B, C and D).

To determine cardiac reversal rate, we created for each fly a square wave trace, where the higher value was attributed to timepoints during forward bouts and the lower value during backward bouts (see Fig. S2 A). We performed Fast Fourier Transform (FFT) on the square waves and found the average cardiac reversal rate to be 0.23 ± 0.02 Hz (mode ± s.e.m., Fig. S2 B) during the baseline period. The mean length of backward and forward bouts during baseline was 1.90 ± 0.04 (mean ± s.e.m) and 1.54 ± 0.05 seconds (mean ± s.e.m), respectively.

Next, we asked whether flies modulate their heart activity when exposed to a visual threat. Since time spent freezing upon looming follows a bimodal distribution (Fig. S1 B), and flies that do not freeze run instead (Fig. S1 C), we separated flies into two groups: *freezers* (n=31) for flies that froze more than 3.5 min and *runners* (n=17) for those that froze less than 1.5 min. Henceforth, we will analyse cardiac activity separately for freezers and runners.

Cardiac arrest during the startle response has been described in all phyla analysed, including invertebrates [9,29], hence we hypothesized that it should be also present in *D. melanogaster* upon a sudden visual threat. Cardiac arrests upon looming should be detectable as a sudden decrease in beating frequency. Hence, we performed a time-frequency analysis of the cardiomyocyte displacement in a window of six seconds around each 500 ms visual stimulus. As expected, we observed that the heart skipped on average one beat during the looming stimulus, reflected in the sudden average decrease 1.35 Hz ± 0.11 (mean ± s.e.m) in beating rate in both freezers and runners (Fig. 2 E). In contrast, we did not observe stimulus driven cardiac arrests in flies exposed to the control stimulus (Fig. 2 E).

It has been previously shown that cardiac arrest is independent of the ensuing behaviour displayed by the animal [9,30]. Similarly, we observed comparable frequency of cardiac arrest upon looming in freezers and runners that was higher than that found at random time points during the baseline period or during the stimulation period at random time points in between looming stimuli *(Mann-Whitney U test,* p<0.0001 and p<0.0001, respectively) (Fig. 2 F). Furthermore, it is known that the startle reflex can be reduced by repetition or anticipation [9]. Indeed, we found that the frequency of cardiac arrest upon repeated looming stimuli decreases over the course of stimulus presentations (55.8% first 4 looming presentations, 39.8% last 4 looming presentations; Wilcoxon test, p= 0.003) (Fig. 2 G), suggesting that in our paradigm the cardiac startle reflex habituates.

Transient cardiac arrests are also observed during cardiac reversal (Fig. 2 A point arrow, and Fig. S2 C) [22,27]. Hence, we asked whether cardiac arrests observed upon looming were associated with cardiac reversal. We found the frequency of arrest associated with looms during which a cardiac reversal was observed to be higher than that observed in looms without cardiac reversal *(Mann-Whitney U test,* p<0.0001; Fig. S2 D). Still, cardiac arrests both with and without reversal were more frequent during looms than during cardiac reversal in between stimulus presentations *(Mann-Whitney U test,* p<0.0001 and p<0.0001; Fig. S2 D). This result indicates that looms trigger cardiac arrests independently of its coupling with a change in pumping direction.

Altogether these results demonstrate that threat leads to an instantaneous cardiac response in the fruit fly, a cardiac arrest, consistent with a startle response, supporting the existence of an autonomic-like reflexive central control of cardiac activity in *Drosophila* that enables quick adaptation to external stimuli.

### Cardiac beating rate decelerates during freezing and accelerates during running

We have shown that looming stimuli trigger a transient cardiac arrest. However, whether exposure to threat leads to long lasting, behaviour specific, modulation of cardiac activity in invertebrates remains unanswered. Thus, we investigated whether the cardiac beating rate is modulated in a behaviour-dependent manner during freezing and fleeing defensive responses.

As mentioned above, the fly heart has different beating frequencies during backward and forward pumping bouts [22,28]. To perform SFT analysis during backward or forward bouts, for each fly we concatenated all backward or forward bouts independently and compared the beating rate during baseline and stimulation periods. Unfortunately, the beating frequency during forward bouts per individual was too variable (Figure 2 D) to allow the observation of reliable changes in heart beating rate during forward pumping mode (Fig. S2).

The distribution of beating frequencies during the backward pumping mode revealed very small, not statistically significant, differences between baseline and looming exposure periods for both freezer and runner flies (Fig. 3 A-C). These small differences were however in opposite directions. Therefore, we compared the change, relative to baseline, in the distribution of backward beating frequencies (stimulation – baseline) across freezers, runners and control flies (Fig. 3 D). We found that relative to control flies, freezer animals increased the prevalence of lower frequencies (around 4 Hz) (Kolmogórov-Smirnov test (K-S), p=0.0008). Conversely, runner flies increased the prevalence of higher frequencies (around 7.5 Hz) (K-S test, p<0.0001). Furthermore, a within animal comparison of backward beating rate before and after stimulation revealed a decrease in the peak rate during stimulation in freezer flies and an increase in both runner and control stimulated flies (Wilcoxon test, p=0.008, p=0.006 and p<0.0001, respectively; Fig. 3 F). The increase in beating frequency in control flies may reflect the weak but reliable increase in walking speed of these flies upon stimulation (Fig. 1 E-F). We observed no differences between freezers, runners and control animals in the peak rate of backward beating before stimulation (one-way ANOVA, p=0.72; Fig. 3 E).

**Figure 3.**
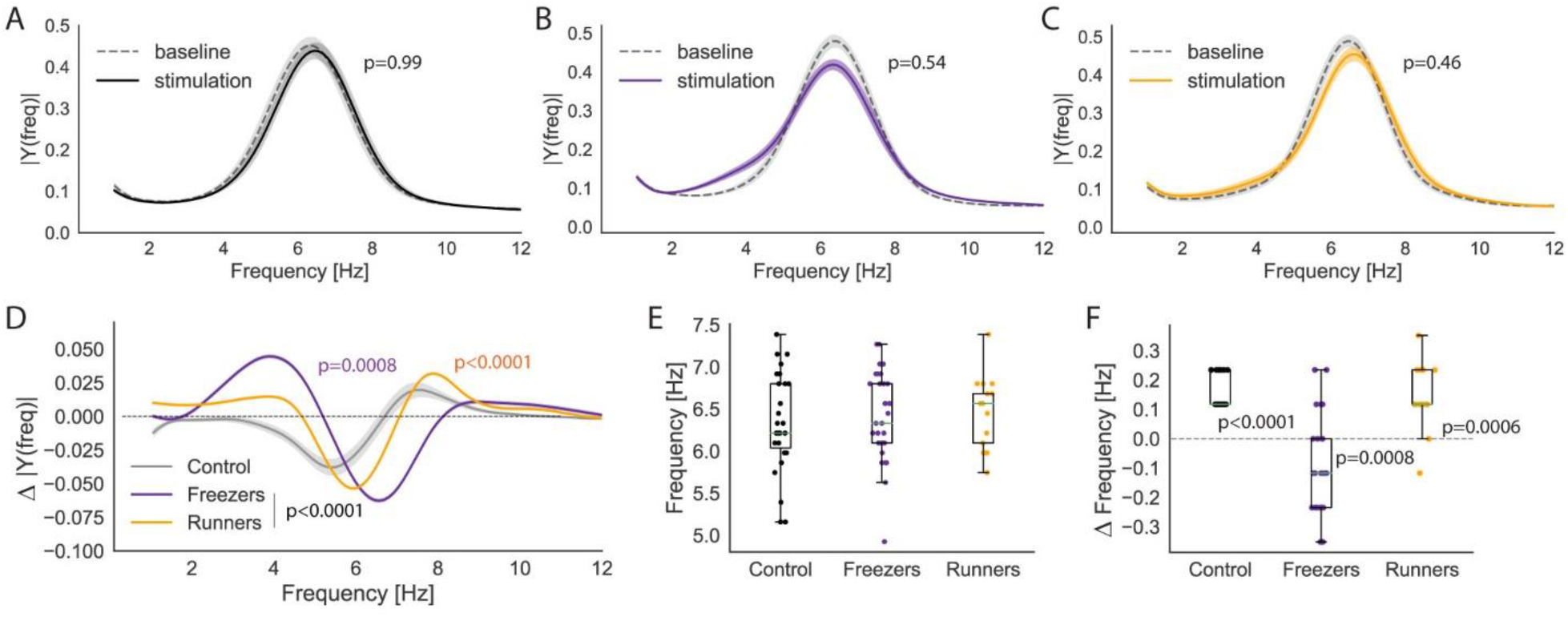
Backward beating rate is regulated in opposite directions while freezing and fleeing. (A-C) Average power spectral density of backward beating rate (± s.e.m.) of control flies (A), freezers (B) and runners (C). (D) Change in the average power spectral density of backward beating rate (± s.e.m.) (stimulation-baseline period). (E) Baseline peak rate of backward beating per individual fly. (F) Change in the peak rate of backward beating per individual fly (stimulation - baseline period). In B-F, black, purple and orange correspond to controls, freezers and runners respectively.

These results show that flies adjust cardiac beating rate during defensive behaviours. While pumping in the backward direction, i.e. towards the abdomen, flies showed bradycardia when freezing and tachycardia when fleeing.

### Cardiac reversal rate decelerates during freezing and accelerates during running

Next, we examined the effect of threat on cardiac reversal. The effects of diverse external and internal stimuli on cardiac reversal have been studied in different species of insects [31,32]. However, the modulation of cardiac reversal upon threat detection remains untested.

To investigate reversal activity, we performed FFT on cardiac reversal signals (Fig. 4 A, B, see also Fig. S2 A) during baseline and during stimulation for bouts of 10 sec or longer of freezing in freezer flies or running in runner flies. To detect habituation or sensitization of the effect of looms on cardiac reversal, we analysed cardiac reversal frequency at early or late phases of the stimulation period. Noticeably, in freezers, freezing was accompanied by a decrease in cardiac reversal rate that was more evident for late freezing bouts when compared to baseline (K-S test, p<0.0001; Fig. 4 C). In contrast, runners increased the reversal rate during fleeing bouts compared to baseline (K-S test, p<0.0001). In this case the increase in reversal frequency was equivalent for early and late running bouts (K-S test, p<0.91) (Fig. 4 E). Interestingly, even before loom presentations, freezers and runners showed a different distribution of reversal frequency (K-S test, p=0.038; Fig S3 A). Freezers showed slower and more variable peak reversal rates *(Mann-Whitney Utest,* p=0.01 and p= 0.0007; Fig. S3 B, C).

**Figure 4.**
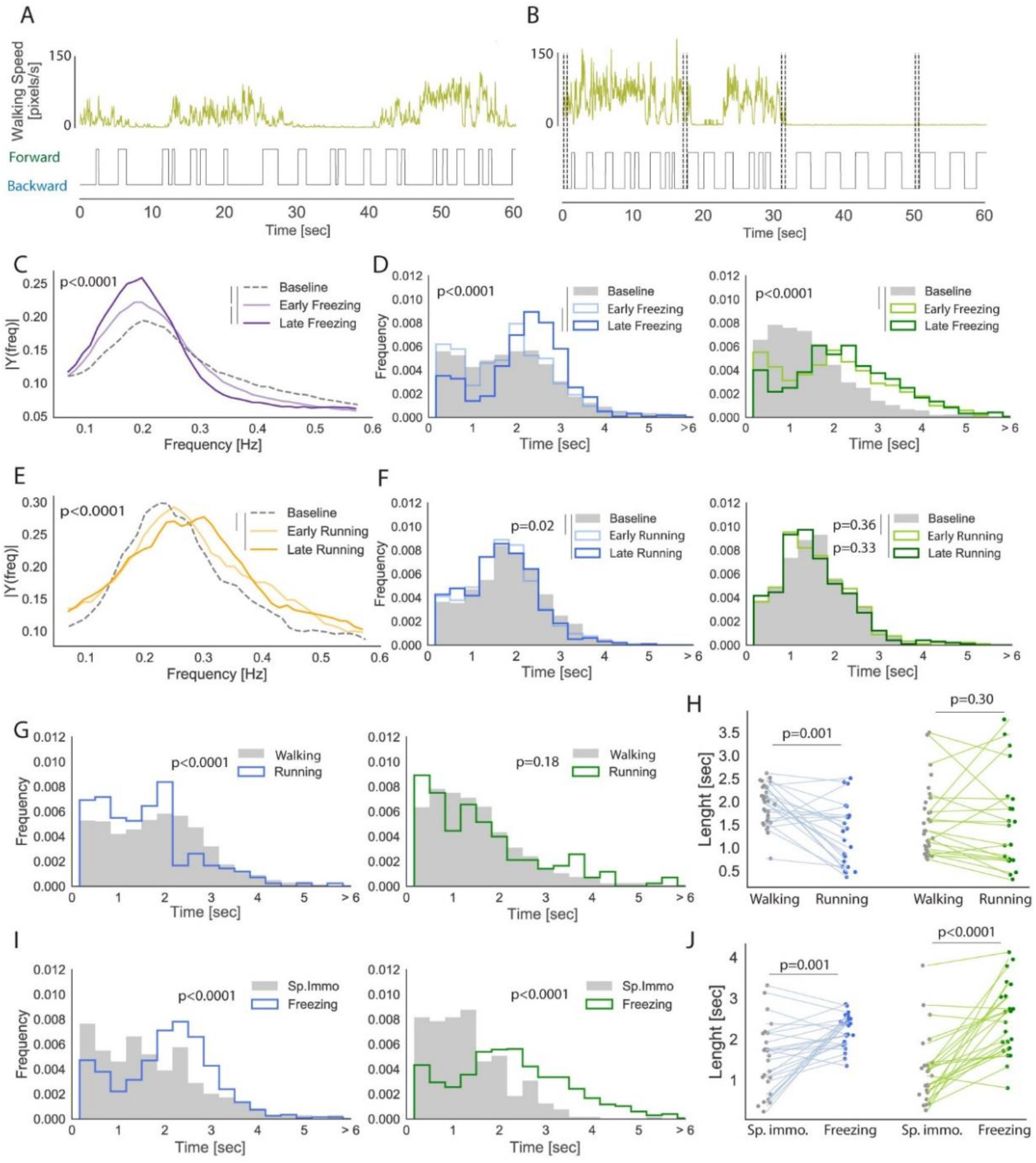
Reversal rate is downregulated during freezing and upregulated during fleeing. (A-B) Representative traces showing walking speed (above) aligned to cardiac reversal (square wave, below) during baseline (A) and stimulation (B). Vertical dashed lines in B represent beginning and end of the stimulus. (C) Power spectral density of cardiac reversal rate during baseline, early and late bouts of freezing for freezers flies. (D) Density distribution of the length of backward (left) and forward bouts (right) of freezers during baseline, early and late periods of freezing during the stimulation period. (E) and (F) Same as C and D for runners during baseline, early and late bouts of running during the stimulation period. (G) Density distribution of the length of backward (left) and forward bouts (right) while walking during baseline and while running during stimulation period for freezers. (H) Median length of backward and forward bouts per individual animal while walking during baseline and running during stimulation, for freezers. (I) Density distribution of the length of backward (left) and forward bouts (right) while spontaneously immobile during baseline and while freezing during stimulation for freezers. (J) Median length of backward and forward bouts per individual animal during spontaneous immobility and freezing for freezers.

Reversal rate and its variability directly depend on the length of the forward and backward pumping bouts. Therefore, we assessed whether cardiac reversal changes during defensive behaviours resulted from a modulation of the length of bouts in backward, forward or both pumping modes, for which we analysed the distribution of bout length for both modes. In agreement with the Fourier analysis of reversal rate, we observed differences in the distribution of both forward and backward bout lengths between freezers and runners during baseline. (Fig. S3 D-E).

During the stimulation period, freezers initially increased the length of forward bouts compared to baseline, and only during late freezing bouts did they also increase the length of backward bouts (K-S test, p<0.0001, p<0.0001 and p<0.0001, respectively; Fig. 4 D). In the case of runners, only backward bouts decreased their length during running periods, a change observed already at the beginning stimulation period that remained stable until the end of stimulation (K-S test, p<0.02 and p<0.02 respectively; Fig 4 F). For a within animal analysis we took the median bout length of forward and backward pumping modes during either freezing, by freezer flies, or running, by runner flies, and compared it to the median bout length during baseline irrespective of the flies’ behaviour. Again, we found that freezers, while freezing, increase the length of both backward and forward bouts *(Mann-Whitney U test,* p=0.001 and p<0.0001, respectively; Fig. S4 A) while runners specifically decreased median backward bout length *(Mann-Whitney U test,* p=0.04 and p=0.46; Fig. S4 B). Importantly, no changes were observed in the length of bouts in control flies (K-S test, p=0.949 and p=0.152 respectively; Fig. S4 D-E). These results suggest that freezing and running are paralleled by the regulation of cardiac reversal in opposite directions: a deceleration of cardiac reversal reflected in increased length of both backward and forward bouts for freezers; and an acceleration of cardiac reversal reflected in decreased length of backward bouts for runners.

These changes may either reflect a trait difference in cardiac modulation between freezers and runners or instead be related to the behaviour expressed. To disambiguate between the two possibilities, we asked whether the same animal could flexibly change its cardiac reversal rate, by decreasing it during freezing and increasing it during fleeing responses. To this end, we focused on freezers animals that mostly freeze in response to looms but showed enough periods of running to perform the analysis of backward and forward bout length during both freezing and running behaviours. In addition, we specifically compared cardiac activity while freezer flies were immobile during stimulation, i.e. freezing, with that when these flies were immobile during baseline, i.e. spontaneous immobility. A similar comparison was made while freezer flies were walking during baseline and running during stimulation. We found that freezers, just like runners, decreased the length of backward bouts while running when compared to the bout length while walking during the baseline period (K-S test, p<0.0001; *Mann-Whitney U test,* p=0.001; Fig. 4 G, H). Indeed, the same animal showed increased bout length while freezing and decreased bout length while fleeing (Fig. S4 C). These data demonstrate that flies flexibly adjust cardiac reversal rate during the expression of distinct defensive behaviours. Furthermore, we observed that the both backward and forward bouts were longer during freezing than during spontaneous immobility (p=0.001 and p<0.0001, respectively; Fig. 4 I, J), consistent with a specific decrease in reversal rate while freezing in response to threat.

All together, these results show that fruit flies flexibly regulate reversal rate upon threat perception in a behaviour specific manner, since flies are bradycardic while freezing and tachycardic while running. Although, it is well known that in vertebrates threat induced freezing is accompanied by bradycardia, whether the latter is part of a specific defensive response or constitutes a generic feature of cardiac activity accompanying any form of immobility remained unanswered. Our findings clearly demonstrate that freezing is a different physiological state from spontaneous immobility.

### Freezing leads to a predominance of forward cardiac pumping and sugar mobilization

We have shown that flies independently modulate the length of backward and forward bouts during threat stimulation. In addition, we found that during freezing the increase in forward bout length is higher than that observed for backward bouts (mean increase in forward bout length was 1.03 ± 0.09 vs 0.27 ± 0.07 seconds for backward bouts; Wilcoxon test, p<0.0001; Fig. S4 C), while during running, flies only decreased the length of backward bouts (Fig. 4 G-H). Hence, the total time the heart beats in forward mode seems to increase upon threat presentation. Thus, we determined the percentage of time in backward pumping, the more reliable pumping mode, during baseline and stimulation periods. The complementary percentage is the proportion of time pumping forward. Before stimulation the heart pumped on average 55.32 ± 0.96% (mean ± s.e.m.) of time in backward mode. No differences in the proportion of time pumping in backward mode were observed between freezer, runner and control flies (one-way ANOVA, p=0.67; Fig. S5 A). Interestingly, during stimulation both freezers and runners, but not controls, decreased the total time pumping in backward mode (Wilcoxon test, p<0.0001, p=0.02 and p=0.09, respectively; Fig. 5 A). The heart of flies pumped in this mode on average 46.29 ± 2.0% (mean ± s.e.m.) of the time in freezers, 51.59 ± 1.49% (mean ± s.e.m.) in runners and 56.84 ± 1.62% (mean ± s.e.m.) in control flies.

**Figure 5.**
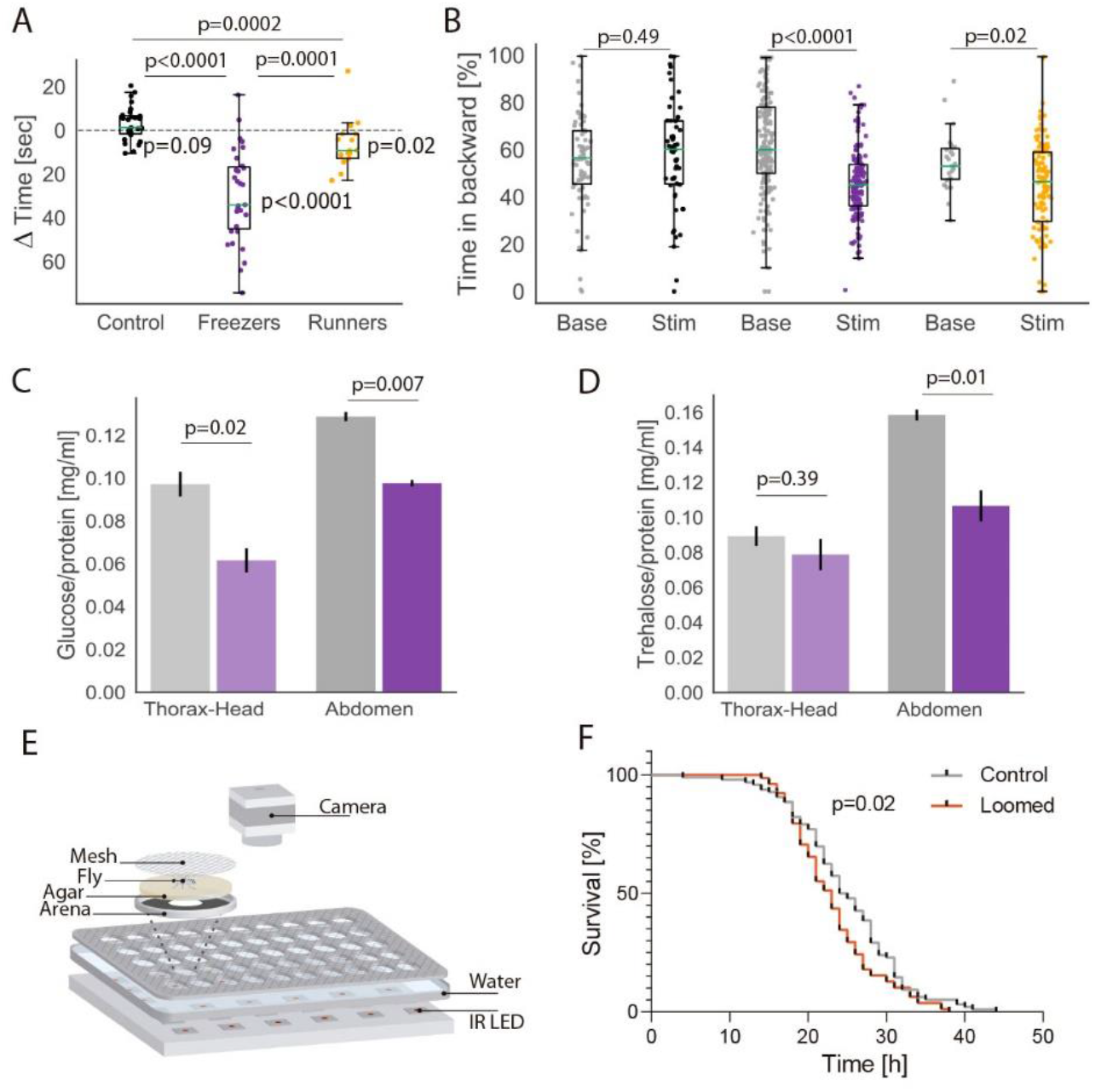
Favoured forward mode and energy mobilization during freezing. (A) Change in total time in backward mode (stimulation – baseline period) for control, freezer and runner flies. (B) Percentage of time in backward mode per period of immobility during baseline (grey) or stimulation in controls (black), freezers (purple) and runners (orange). (C-D) Glucose (C) and trehalose (D) concentration in the hemolymph (Thorax or Abdomen) from control (grey, n=2 replicates with a total of 20 flies) and freezer (purple, n=3 replicates with a total of 30 flies) flies. Mean (± s.e.m.) across 2 and 3 replicates respectively are shown. (E) Schematic representation of the starvation setup. (F) Resistance to starvation of flies after 15 min of stimulation (n=2 replicates for a total of 98 control flies and 98 loomed flies).

The stronger decrease in the percent time pumping in backward mode observed in freezers may result from the difference in behaviour of freezers and runners, since the former spent most of the stimulation period immobile and the latter spend most of the time running. To address this issue, we determined the percent time pumping in backward mode exclusively during bouts of immobility (both spontaneous immobility bouts during baseline or looming triggered freezing) for freezers, runners and control animals. We found that both freezers and runners, while freezing, decreased the total time spent in backward pumping mode even when compared with spontaneous immobility during baseline *(Mann-Whitney U test,* p<0.0001 and p=0.02; Fig. 5 B). In contrast, control flies showed no such decrease during immobility bouts in the stimulation period *(Mann-Whitney U test,* p=0.49; Fig. 5 B). The observed cardiac modulation during freezing led the heart pumping in backward mode on average 45.44 ± 1.04% (mean ± s.e.m.) of the time. We found no differences in the percentage of time in backward pumping during walking/running periods in freezer, runner, or control flies (Fig. S5 B). These findings show that flies favour forward pumping, i.e. towards the thorax and head, specifically during freezing and again that this defensive behaviour corresponds to a different internal state from that found during spontaneous immobility.

Backward and forward pumping of the heart promote the flow of the hemolymph ensuring the transport of nutrients, metabolic waste and hormones between the thorax-head and abdomen. Hence, we hypothesized that the observed increase in forward pumping mode might reflect an increased flux of nutrients from the fat body, the fly’s main energy store located in the abdomen, to the head and thorax to meet the increased energy demands while freezing. We focused on carbohydrates, namely glucose and trehalose, as these are an important source of energy. Ingested glucose can be taken up by any tissue and be quickly metabolized providing energy to the cells or be used to create energy stores in the form of glycogen [33]. Trehalose, a disaccharide composed of two α-glucose molecules, is synthetized from glycogen in the fat body and then secreted into the hemolymph, constituting the major circulating sugar in flies and a crucial source of energy to the brain and muscles [33]. We measured glucose and trehalose levels in the hemolymph separately from abdomens and thorax-head of flies after prolonged freezing and compared to control flies. To maximize freezing behaviour, we designed small arenas (1,6 cm Ø), as flies are more likely to freeze in small confined spaces, and exposed flies to 60 looms or control stimuli over the course of 15 minutes. In these conditions, loom-stimulated animals froze 96.53 ± 3.53% (mean ± s.e.m.) of the time during the stimulation period while control-stimulated flies were immobile for 36.6 ± 0.78% (Fig. S5 C-F). We found that glucose levels were decreased in both thorax-head and abdomen in loom-stimulated flies compared to control-stimulated (t-test; p=0.02 and p=0.007; Fig. 5 C). Strikingly, we found trehalose levels to be reduced in the abdomen but not in the thorax and head (t-test; p=0.01 and p=0.39; Fig. 5 D), which is in line with our finding that the heart increases pumping towards these body compartments.

These findings show that loomed flies reduce sugar levels in the hemolymph. Although a decrease in trehalose suggests an increase in energy consumption, a decrease in glucose could result from either an increase in energy consumption or an increase in energy stores through glycogen biosynthesis [33]. Since the rate of energy consumption directly impacts starvation resistance [34], next we tested if this was altered in flies after prolonged freezing compared to control flies. We stimulated flies in enclosed arenas over 15 min with 60 looms or control stimuli, during which loomed flies froze the entire time (Fig S5 C-F), and then recorded their activity during the next four days. To ensure access to water, we filled the bottom part of each arena with agar 1% and maintained the humidity after stimulation (see methods and Fig. 5 E). We analysed the movement of each fly and annotated their death as occurring when no pixel change was observed for 1 h. We found loomed flies to be less resistant to starvation than control stimulated flies (Log-Rank test, p=0.02; Hazard ratio=1.47, 95% CI= 1.054-2.04; Fig. 5 F).

Together these data demonstrate that freezing behaviour is costly, as it renders flies less resistant to subsequent starvation, likely due to an increase in the consumption and mobilization of sugars. Our data although consistent with the view of freezing behaviour as a form of detection avoidance while preparing the animal for future action, does not support the view of freezing as an energy sparing state.

### Cardiac reversal rate and variability predict threat-induced freezing

Having shown that cardiac reversal regulation is a physiological feature of defensive behaviours in *Drosophila* and that the pattern of reversal is different even in resting conditions for freezers and runners, we subsequently asked whether cardiac activity predicts the behavioural response to threat. Cardiac reversal is slower and more variable in flies that responded to looming mostly by freezing than that of flies that responded by running *(Mann-Whitney U test,* p=0.01 and p= 0.0007; Fig. S3 B, C). Since these are independent variables (Pearson=0.01, p=0.92; Fig. S6 A), we modelled the amount of time flies spent freezing upon looming using ordinary least squares (OLS) with peak reversal rate and reversal rate variability during baseline as predictors. We found that our model explained 29.2% of the variance of time the flies spend freezing (F-stat<0.0001; Fig 6 A).

**Figure 6.**
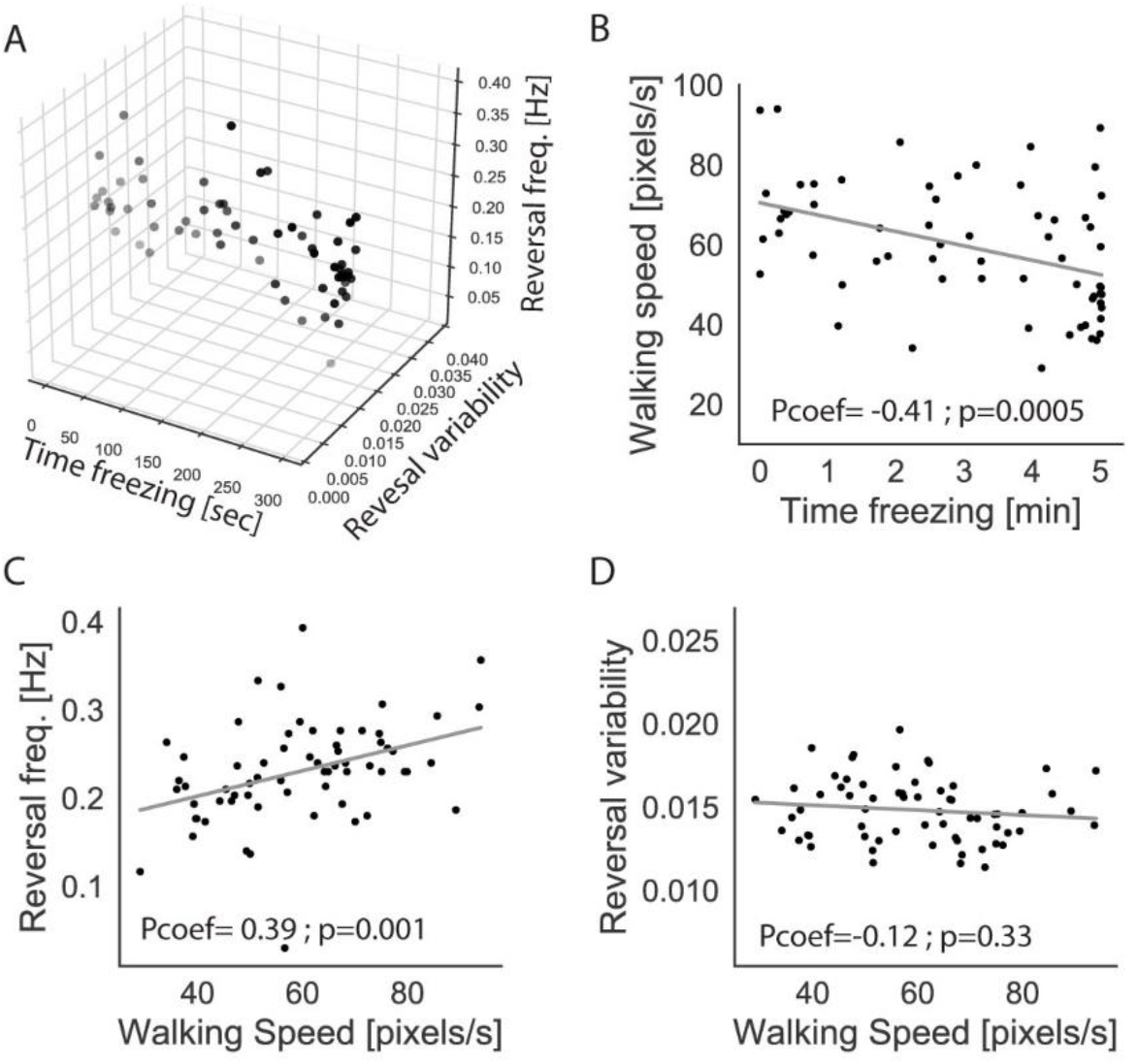
Cardiac reversal rate and variability predict freezing response. (A) 3D scatter plot showing the total time freezing, peak cardiac reversal rate and variability for the 64 loomed flies. (B-D) Scatter plot showing the value of baseline average walking speed and the total time freezing (B), reversal frequency (C) and reversal variability (D) for each loom stimulated fly. Least squares regression lines are shown in grey. Pcoef shows Pearson coefficient.

Interestingly, these results demonstrate that cardiac activity in flies predicts the defensive response upon looming. However, baseline differences in cardiac reversal between freezers and runners might reflect differences in behaviour before stimulation. In our experimental conditions, both walking speed and total time spent walking during the baseline period correlated with total time freezing upon loom stimulation (Pcoef=-0.41; p=0.0005 and Pcoef=-0.37; p=0.02, respectively; Fig. 6 B and S6 B). Since walking speed and time spent walking were correlated (Pcoef=0.58; p<0.0001; Fig. S6 C), henceforth we will use walking speed as a variable to measure locomotor activity of the fly during baseline. We found that walking speed is correlated with peak cardiac reversal rate (Pearson=0.39; p=0.001; Fig. 6 C). However, no correlation was found between baseline reversal rate variability and walking speed (Pearson=-0.12; p=0.33; Fig. 6 D). These results suggest that cardiac reversal rate, but not cardiac reversal variability, is dependent on ongoing behaviour. Therefore, whether the relevant predictor of time spent freezing during exposure to looming stimuli is cardiac reversal rate or walking speed during the baseline, remains to be established. Indeed, when we tested walking speed, instead of cardiac reversal rate, together with cardiac reversal variability as predictors of time spent freezing using OLS, we found that once more they explained 32,4 % of the variance between animals (F-stat<0.0001) (Fig. S6 D).

Altogether, these results show that two independent features of cardiac reversal, rate and variability, are predictors of future levels of looming triggered freezing. We show that cardiac reversal variability is independent of ongoing levels of locomotor activity suggesting that it might reflect a trait of the animal that will impact the defensive response.

## Discussion

### Functional role of freezing coupled bradycardia

The prevalence of fleeing and freezing responses across vertebrate and invertebrate groups raises the possibility that the physiological responses observed in vertebrates are also present in invertebrates. Our data demonstrate for the first time that *D. melanogaster* regulates its cardiac activity upon threat perception. Although homologous human and fly genes have been implicated in cardiomyocyte specification, the functional architecture of the fly’s heart is most likely analogous to the chambered heart of humans. In addition, the neuronal control of cardiac reversal, absent in vertebrates, is likely to constitute an independently evolved mechanism mediating cardiac function regulation. Still, the fruit fly downregulates cardiac output (cardiac beating and cardiac reversal) when freezing and upregulates it when fleeing, just like vertebrates do. Further, it has been demonstrated in other insects [19–21], that cardiac reversal regulation leads to changes in tracheal ventilation, the latter might also accelerate during fleeing and decelerate during freezing.

Several hypotheses have been put forth to explain the characteristic bradycardia that accompanies defensive freezing behaviour. The motivational approach proposes that the downregulation of cardiac activity during freezing might meet the need for decreased energy consumption as part of the preparation for future action [35] Indeed, in frugivorous bats cyclic bradycardia while resting allows the reduction of the amount of energy used per day [36]. The cognitive model proposes that cardiac deceleration may help the brain increase sensory processing [37]. A third hypothesis proposes that bradycardia was inherited from our vertebrate aquatic ancestors that show bradycardia in the context of threat-induced diving responses, where lowering oxygen consumption is crucial [10]. Our findings demonstrate that downregulation of cardiac activity during freezing is already present in invertebrates and suggests that its adaptative value is not related to a decrease in the metabolic needs. In this work, we show that despite the observed bradycardia when flies freeze upon looming, this is a costly behavioural state. Whether freezing is energetically costly in vertebrates remains to be tested.

### Flexible control of cardiac activity

In vertebrates, modulation of cardiac activity that accompanies defensive behaviours is mediated by the autonomic nervous system. Flight-or-fight responses are associated with stronger sympathetic activity that results in tachycardia [4-6]. By contrast, freezing response leads to parasympathetically mediated bradycardia [1,7,8]. It has been argued that although there is no structural counterpart to the autonomous nervous system in invertebrates, the basic functional properties may have been established very early in metazoan evolution [38,39]. The heart of fruit fly is innervated at least by two neuronal populations: the transverse nerves and the bipolar neurons, immunoreactive for glutamate and CCAP respectively [40]. These were shown to regulate cardiac beating and cardiac reversal [28,41] and may act as two independent pacemakers [40]. The independent neuronal control of backward and forward pumping is in line with our findings revealing independent regulation of the length of backward and forward bouts during defensive responses. Furthermore, we show that upon threat detection flies quickly regulate their cardiac activity supporting the idea of an autonomic-like reflexive control of cardiac activity in *Drosophila.* Further studies are needed to explore the existence of quick neuronal control of cardiac activity during defensive behaviours.

### Cardiac activity as modulator of freezing

It is well known that the decision to freeze depends on several factors from the external and internal environment [23,24,42–45]. In flies, we have shown that the animal’s own behavioural state [11] and the social environment regulates threat-induced freezing [15]. Here, we show that two features of cardiac activity measured before threat, reversal rate and reversal variability, partially predict the amount of looming triggered freezing. Furthermore, we observed that reversal rate correlates with ongoing locomotor activity, widely used as a proxy for arousal in all taxa including flies [46], suggesting that cardiac reversal rate might be an additional measure of physiological arousal. Further studies on the relationship between cardiac reversal rate, locomotion and arousing stimuli will be instrumental for our understanding of physiological arousal and its relevance to behaviour.

In contrast, we found no correlation between cardiac reversal variability and the ongoing locomotor behaviour before stimulation. Whether cardiac variability reflects a transient internal state of the animal or a persistent individual trait remains to be established. Interestingly, heart rate and heart rate variability are used in humans as proxies for the status of the parasympathetic and sympathetic nervous system and as predictors of health and emotional responses [47,48]. To definitively establish cardiac activity as a modulator of defensive behaviours, a mechanistic understanding of cardiac regulation it is required.

### Summary

This study shows the prevalent coupling between defensive responses and cardiac function in an invertebrate species. It reveals that flies regulate cardiac pumping (backward and forward) in a fast, flexible and independent manner, suggesting independent neuronal modulators that may resemble some features of the vertebrate autonomous nervous system. In addition, it demonstrates that threat-induced freezing corresponds to an internal state distinct from that found during spontaneous immobility. Finally, it establishes that energy sparing and inheritance of traits from aquatic vertebrates are unlikely explanations for bradycardia during freezing behaviour, favouring a possible role in sensory processing. This is in accordance with recent literature showing in humans that the brain processes information regarding cardiac function and that this affects processing of sensory information and memory formation [49–51]. We believe this work establishes the fruit fly as a model well suited to mechanistic, functional and comparative studies on the regulation of cardiac physiology and its contribution to behaviour.

## Materials and Methods

### Fly lines and husbandry

To image cardiac activity, we used 1-2 days old mated females fruit flies (*Drosophila melanogaster*). The following stock was used: yw^-^; *tinCGal4,* UAS-nls-dsRed/CyO. *tinC*Gal4 gift from Manfred Frasch [52], UAS-nls-dsRed (Bloomington Drosophila Stock Center, #8546). Flies were reared on standard food at 25° C and 70% humidity in a 12 h-12 h dark:light cycle.. For aging, 20 females and 5 males were transferred to a fresh vial for 24 h.

For starvation resistance and sugar measurements experiments 10 to 11 days-old mated CantonS males were raised at 25° C and 70% humidity in a 12 h-12 h dark:light cycle. For aging, 15 females and 15 males were transferred to a fresh vial every 72 h.

### Handling of flies

#### Tethering procedure

Flies were briefly cold anaesthetised and glued to a coverslip using Norland Optical adhesive #61 cured with a 365 nm UV lamp (UVP) for 30 s. Each fly was given 10 min to recover before starting an experiment. In total, 35 flies were dots (control) stimulated and 80 were loomed. From those only 27 and 64 flies respectively had sufficient cardiac fluorescence signal to be analysed. All behavioural experiments were performed in the 8 hours period after light onset.

#### Testing arenas

Flies were cold anaesthetised for 10 min before being placed into the arenas. Before starting the experiment, flies were given 15 min to recover. In total, 49 control and 49 loom stimulated flies were used per experiment. All experiments were performed in the 8 hours period after light onset.

### Visual stimulation

After a 5 minutes baseline period, experimental and control animals received 20 (heart imaging setup) or 60 (testing arenas) stimuli presentations with an ISI ± 20 seconds.

All visual stimuli in the heart imaging set-up were generated as described in Zacarias et al. (2018) [11] in custom python scripts using PsychoPy [53]. To generate a looming effect, a black circle increased in size on a blue background. The visual angle of the expanding circle was determined by the equation: *θ(t)*= 2tan^-1^ (*l*/ *vt)* (Eq. 1), where *l* is half of the length of the object and *v* the speed of the object towards the fly. The virtual object length was 1 cm and the speed was 25 cm s^-1^ (*l*/ *v* value of 40 ms). Each looming presentation lasted for 500 ms. The object expanded for 450 ms until it reached maximum size of 78° where it remained for 50 ms before disappearing. Synchronous with expansion, the looming stimulus produced a considerable decrease in luminance within the behavioural apparatus. We measured luminance using a digital lux meter (DX-100, INS instrumentation). When no stimulus was being presented (blue screen) the luminance at the stage was 260 lux. Just before looming offset, when the disk reached its maximum size, luminance was 32 lux, representing an 88% decrease. To control for these changes, we created a stimulus where an array of approximately 5° dots was added each frame in random positions as not to create an expanding pattern. The size and number of dots was determined empirically to generate a similar decrease in luminance as the looming stimulus (35 lux, 86.5% decrease).

For stimulation in enclosed arenas, similar visual stimuli were generated using a custom Bonsai workflow.

### Behavioral apparatus

#### Semi-restrained flies

Cardiac activity was imaged from semi-restrained flies using a Hamamatsu C11440 camera (ORCA-flash4.0LT) and a 10X/0.30w objective (Olympus). Excitation light was provided by a 530 nm mounted LED from Thorlabs. A small porexpam ball (4.3 mm diameter) was held by a custom-made holder (the funnel section of a glass Pasteur pipette) underneath the fly at a distance that allowed the fly to catch the ball. The ball could be freely moved by the fly and if released, the fly could catch the ball again. Visual stimuli were presented on a 24-inch monitor (ASUS VG248QE) with a 144 Hz refresh rate and tilted at 45 degrees over the imaging stage. To visualize the presentation of stimuli in the recorded video, we used a programmable microcontroller (Arduino Mega 2000) to trigger the illumination of a custom-made red LED synchronized with the stimulus presentation. To image the fly’s movement, an infrared (850nm) LED was placed laterally to illuminate the fly. Fly behaviour and ball movement was recorded using a USB3 video camera (Flea3 1.3 MP Mono USB3 Vision, Point Grey, Richmond, Canada) with a 700 nm long pass filter placed 40 cm away from the imaging stage. The entire rig was placed inside a light-tight black enclosure.

#### Testing arenas

Two identical behavioural apparatus were used in parallel, each one with a stimulation monitor (Asus ROG Strix XG258Q, 24.5”) tilted at 45° over the stage running at 240 Hz refresh rate. 50 circular testing arenas (16 mm in diameter) were cut in white acrylic 4 mm thick plates. For the starvation resistance assay, 3 of these plates were stacked together such that each arena was 12 mm high, of which 8mm were filled with agar 1%, leaving 4 mm for the fly to move. To prevent flies from escaping and allow air exchange and video acquisition the plates were covered with nylon socks. For sugar measurement experiments and scoring of freezing behaviour, the arenas were made of one acrylic plate (such that each arena was 4 mm high) on top of a 3 mm thick, white opalino sheet that allowed diffuse light through. The plate was covered with a transparent acrylic sheet for maximum resolution of the fly movement throughout the stimulation session. To increase the resolution of the image for subsequent behaviour quantification only 15 flies were recorded. We used a USB3 camera (FLIR Blackfly S, Mono, 1.3MP) at 60 Hz with a 730 nm long pass filter (Lee Filters, Polyester 87 Infrared). The stage plates were backlit by an infrared (940 nm) LED array developed by the Scientific Hardware Platform at the Champalimaud Centre for the Unknown. The entire rig was kept at 25° C and 70% humidity.

#### Starvation resistance

Immediately after stimulation the two stage plates were placed in a transparent container with water, ensuring that the agar is moist until the end of the experiment. The container was backlit by a custom-built infrared (940 nm) LED array, and a camera (the same as in the stimulation setup) recorded both stage plates at 6 Hz. The entire rig was kept at 25° C and 70% humidity in a 12 h-12 h dark-light cycle until the last fly died.

### Video acquisition and tracking

#### Semi-restrained flies

Fly videos were acquired using Bonsai [25] at 60 Hz and width 1104 x height 1040 resolution. We extracted two main features from the videos using Bonsai [25]: ball movement and fly motion. Walking speed was extracted from the displacement in 3 dimensions of feature points in the surface of the ball. Fly motion was quantified by the number of pixels active in a 162×117 pixel region of interest surrounding the fly (ROI, see supplementary 1). A minimum change of 18 intensity levels from one frame to the next was required for a single pixel to be considered active.

Heart videos were recorded using HC Image Live Software at 30 Hz obtaining a standard 8bit RGB tiff file as the signal. Tracking of cardiomyocytes nuclei was performed using Bonsai [25]. The displacement perpendicular to the heart axis was represented by the x coordinate in a ROI surrounding the nucleus.

#### Testing arenas

We used a custom-built Bonsai workflow [25] for video acquisition at 60 Hz and to extract to main features of the fly: centroid position and pixel change in a 72 x 72 pixels ROI around the fly.

#### Starvation resistance

We used a custom-built Bonsai workflow [25] for video acquisition at 1 Hz and to quantify pixel change inside each arena. A pixel was scored to be active if it recorded a change higher than 5 values of intensity from the previous frame.

### Glucose and trehalose assays

Immediately after visual stimulation, plates with 50 flies were placed on dry ice. We collected 10 thorax-heads or abdomens per sample. Dissected body parts were rinsed twice with cold PBS and then homogenized in 200 μl of cold PBS. We spun down samples for 1 min at 5000 rpm at 4° C. After keeping aside 15 μl for the assay (Pierce BCA protein Assay Kit, #10678484, Fisher Scientific), the supernatant was heated for 5 min at 70° C. Next, we spun down for 3 min at 13.000 rpm at 4° C and performed glucose assays on the supernatant using a Glucose (GO) assay kit from Sigma (GAGO-20).

For the trehalose assays we incubated samples overnight with trehalase (#T0167, Sigma). Absorbance was measured in a plate reader (BMG Labtech, SpectroStar Nano) at 562 nm for protein measurement and 540 nm for glucose. Two and three samples were measured for control and loom stimulated animals.

### Data analysis from semi-restrained flies

Data were analysed using custom scripts in Python 2.

#### Behavioural classifiers

To classify behavioural states, walking speed and fly movement data were averaged over a 83.3 ms (5 frames) moving window to smooth out short-term fluctuations. Walking speed was determined from data extracted from the ball movement, using the following formula: 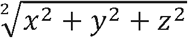. The threshold for classifying not walking periods was determined by manual annotation. Non-walking periods were classified as periods longer than 2.5 sec with a walking speed lower than 0.5 pixels/frame. Fly activity was extracted from pixel change data. The threshold for classifying not moving periods was determined by manual annotation. Immobility/freezing was classified as periods longer than 2.5 sec periods with a pixel change lower than 5 pixels/frame within the ROI.

#### Backward/Forward modes

We regularized datasets of the displacement of the nuclei of cardiomyocytes by subtracting the mean and dividing by the standard deviation. The heart signal was up-sampled to 60 Hz using linear interpolation to reach the same sampling that we had from fly videos. We determined backward bouts by peak to peak comparison between the anterior and posterior nucleus traces. A contraction was classified as backward pumping only if an anterior cell peak happened either at the same time or followed by a posterior cell peak in the next 7 frames. If two backward contractions were separated by more than 18 frames we treated them as two different bouts, and we classified as a forward bout the contractions in between. Only bouts of more than 3 backward contractions were included. We assumed that the intervals between two backward bouts were forward bouts.

#### Cardiac arrest

A cardiac arrest was classified as a sudden decrease in cardiac beating frequency between two consecutive frames above 0.2 Hz. Cardiac arrest was associated with looming when it was found in a 666.6 ms window including each looming. To determine the frequency of cardiac arrests outside looming presentation, we performed a randomization test (with 1000 shuffles with 20 random points per animal) during baseline or stimulation period excluding looming windows.

#### Cardiac reversal

Cardiac reversal frequency was determined by performing a Fast Fourier Transform on the square waves obtained from the analysis of backward and forward bouts. The minimum value was assigned to backward pumping and the maximum value to forward pumping. For analysis of cardiac reversal frequency during different behaviours, we concatenated the square waves for periods longer than 10 sec in which the animal was showing a specific behaviour. Fast Fourier Transform was performed on the square trace resulting from each behaviour including all animals. To analyse early and late phases of freezing, we included the first and last third of freezing bouts, respectively.

For analysis of length of backward and forward bouts during immobility/freezing, only bouts in which the average pixel change was less or equal to 5 were included. Backward and forward bouts in which the average pixel change was above 5 were included as moving periods (walking/running).

### Data analysis from testing arenas

Data were analysed using custom scripts on Python 3. Using the centroid position, a fly was considered to be walking if its speed was higher than 4 mm/s and lower than 75 mm/s. Freezing was classified as the absence of pixel change for periods longer than half a second.

### Data analysis of starvation resistance

Data were analysed using custom scripts on Python 3. Using an hour length step, a fly was considered to be dead if during the last hour less than 3 minutes of motion was recorded. Survival curves were analyzed using Graphpad Prism 8.4.0.

### Data analysis from glucose/trehalose assays

Data were analysed using custom scripts on Python 2.

### Statistics

Prior to statistical testing, the data were tested for normality with a Shapiro-Wilk test and the appropriate non-parametric test was chosen if the data were not normally distributed. All statistical tests are specified in the results section of the text or figure captions and are two-sided. To test whether two samples are drawn from the same distribution we computed a Kolmogorov-Smirnov statistic. Probabilities were compared using the χ^2^ contingency test. Linear regression algorithm was calculated using a simple ordinary least squares model in Python. Survival curves were compared using Log-rank (Mantel-Cox) test. We use sequential Bonferroni correction when multiple comparisons were done to control the familywise error rate.

## References

1. Hagenaars, M.A., Oitzl, M., and Roelofs, K. (2014). Updating freeze: Aligning animal and human research. Neurosci. Biobehav. Rev. 47, 165–176. Available at: http://dx.doi.org/10.1016/j.neubiorev.2014.07.021.

2. Miki, K., and Yoshimoto, M. (2010). Role of differential changes in sympathetic nerve activity in the preparatory adjustments of cardiovascular functions during freezing behaviour in rats. Exp. Physiol. 95, 56–60.

3. Kapp, B. S., Gallagher, M., Applegate, C. D., & Frysinger, R. C. (1982). The amygdala central nucleus: Contributions to conditioned cardiovascular responding during aversive Pavlovian conditioning in the rabbit. In Conditioning (pp. 581–599). Springer, Boston, MA.

4. Cannon, B. (1929). Reviews 1929. Physiol. Rev. IX, 399–431. Available at: https://doi.org/10.1152/physrev.1929.9.3.399.

5. Schenberg, L.C., Corral Vasquez, E., and Barcellos da Costa, M. (1993). Cardiac baroreflex dynamics during the defence reaction in freely moving rats. Brain Res. 621, 50–58.

6. Carrive, P., Bandler, R., and Dampney, R. A. L. Anatomical evidence that hypertension associated with the defence reaction in the cat is mediated by a direct projection from a restricted portion of the midbrain periaqueductal grey to the subretrofacial nucleus of the medulla. Brain research, 460(2), 339–345 (1988).

7. Steen, J.B., Gabrielsen, G.W., and Kanwisher, J.W. (1988). Physiological aspects of freezing behaviour in willow ptarmigan hens. Acta Physiol. Scand. 134, 299–304.

8. Koba, S., Inoue, R., and Watanabe, T. (2016). Role played by periaqueductal gray neurons in parasympathetically mediated fear bradycardia in conscious rats. Physiol. Rep. 4, 1–13.

9. Cuadras, J. (1981). Behavioral determinants of severe cardiac inhibition. Physiol. Psychol. 9, 384–392.

10. Campbell B.A., Wood G., McBride T. Origins of orienting and defensive responses: An evolutionary perspective. In: Lang PJ, Simons RF, Balaban MT, editors. Attention and orienting: Sensory and motivational processes. Hillsdale, NJ: Lawrence Erlbaum Associates; 1997. pp. 41–67.

11. Zacarias, R., Namiki, S., Card, G.M., Vasconcelos, M.L., and Moita, M.A. (2018). Speed dependent descending control of freezing behavior in Drosophila melanogaster. Nat. Commun. 9, 1–11.

12. Takanashi, T., Fukaya, M., Nakamuta, K., Skals, N., and Nishino, H. (2016). Substrate vibrations mediate behavioral responses via femoral chordotonal organs in a cerambycid beetle. Zool. Lett. 2, 19–23. Available at: http://dx.doi.org/10.1186/s40851-016-0053-4.

13. Card, G.M. (2012). Escape behaviors in insects. Curr. Opin. Neurobiol. 22, 180–186. Available at: http://dx.doi.org/10.1016/j.conb.2011.12.009.

14. Gibson, W.T., Gonzalez, C.R., Fernandez, C., Ramasamy, L., Tabachnik, T., Du, R.R., Felsen, P.D., Maire, M.R., Perona, P., and Anderson, D.J. (2015). Behavioral responses to a repetitive visual threat stimulus express a persistent state of defensive arousal in drosophila. Curr. Biol. 25, 1401–1415. Available at: http://dx.doi.org/10.1016/j.cub.2015.03.058.

15. Ferreira, C.H., and Moita, M.A. (2020). Behavioral and neuronal underpinnings of safety in numbers in fruit flies. Nat. Commun. 11. Available at: http://dx.doi.org/10.1038/s41467-020-17856-4.

16. De Vries, S.E.J., and Clandinin, T.R. (2012). Loom-sensitive neurons link computation to action in the Drosophila visual system. Curr. Biol. 22, 353–362. Available at: http://dx.doi.org/10.1016/j.cub.2012.01.007.

17. Anderson, D.J., and Adolphs, R. (2014). A framework for studying emotions across species. Cell 157, 187–200. Available at: http://dx.doi.org/10.1016/j.cell.2014.03.003.

18. Rizki T.M. In: The Circulatory System and Associated Cells and Tissues. Ashburner M., Wright T.R., editors. Academic Press; Waltham, MA, USA: 1978. pp. 397–452.

19. Wasserthal, L.T. (2012). Influence of periodic heartbeat reversal and abdominal movements on hemocoelic and tracheal pressure in resting blowflies Calliphom vicina. J. Exp. Biol. 215, 362–373.

20. Wasserthal, L.T. (2014). Periodic heartbeat reversals cause cardiogenic inspiration and expiration with coupled spiracle leakage in resting blowflies, Calliphora vicina. J. Exp. Biol. 217, 1543–1554.

21. Wasserthal, L.T. (1981). Oscillating haemolymph “circulation” and discontinuous tracheal ventilation in the giant silk moth Attacus atlas L. J. Comp. Physiol. ? B 145, 1–15.

22. Wasserthal, L.T. (2007). Drosophila flies combine periodic heartbeat reversal with a circulation in the anterior body mediated by a newly discovered anterior pair of ostial valves and “venous” channels. J. Exp. Biol. 210, 3707–3719.

23. Scarano, F., and Tomsic, D. (2014). Escape response of the crab Neohelice to computer generated looming and translational visual danger stimuli. J. Physiol. Paris 108, 141–147. Available at: http://dx.doi.org/10.1016/j.jphysparis.2014.08.002.

24. De Franceschi, G., Vivattanasarn, T., Saleem, A.B., and Solomon, S.G. (2016). Vision Guides Selection of Freeze or Flight Defense Strategies in Mice. Curr. Biol. 26, 2150–2154. Available at: http://dx.doi.org/10.1016/j.cub.2016.06.006.

25. Lopes, G., Bonacchi, N., Frazão, J., Neto, J.P., Atallah, B. V., Soares, S., Moreira, L., Matias, S., Itskov, P.M., Correia, P.A., et al. (2015). Bonsai: An event-based framework for processing and controlling data streams. Front. Neuroinform. 9, 1–14.

26. Ocorr, K., Vogler, G., and Bodmer, R. (2014). Methods to assess Drosophila heart development, function and aging. Methods 68, 265–72. Available at: http://www.pubmedcentral.nih.gov/articlerender.fcgi?artid=4058868&tool=pmcentrez&rendertype=abstract.

27. Klassen, M.P., Peters, C.J., Zhou, S., Williams, H.H., Jan, L.Y., and Jan, Y.N. (2017). Age-dependent diastolic heart failure in an in vivo Drosophila model. Elife 6, 1–22.

28. Dulcis, D., and Levine, R.B. (2005). Glutamatergic innervation of the heart initiates retrograde contractions in adult Drosophila melanogaster. J. Neurosci. 25, 271–80. Available at: http://www.ncbi.nlm.nih.gov/pubmed/15647470.

29. Casto, R., Nguyen, T., and Printz, M.P. (1989). Characterization of cardiovascular and behavioral responses to alerting stimuli in rats. Am. J. Physiol. - Regul. Integr. Comp. Physiol. 256.

30. Cuadras, J. (1980). Cardiac responses to visual detection of movement, mechanostimulation and cheliped imposed movement in hermit crabs. Comp. Biochem. Physiol. -- Part A Physiol. 66, 113–117.

31. Ichikawa, T., Ito, K., Ichikawa, T., and Ito, K. Calling Behavior Modulates Heartbeat Reversal Rhythm in the Silkmoth Bombyx mori Calling Behavior Modulates Heartbeat Reversal Rhythm in the Silkmoth Bombyx mori. 16, 203–209.

32. Thon, B. (1980). Habituation of cardiac and motor responses to a moving visual stimulus in the blowfly (Calliphora vomitoria). J. Comp. Physiol. Psychol. 94, 886–893.

33. Mattila, J., and Hietakangas, V. (2017). Regulation of carbohydrate energy metabolism in Drosophila melanogaster. Genetics 207, 1231–1253.

34. Rion, S., and Kawecki, T.J. (2007). Evolutionary biology of starvation resistance: What we have learned from Drosophila. J. Evol. Biol. 20, 1655–1664.

35. Vila, J., Guerra, P., Muñoz, M.Á., Vico, C., Viedma-del Jesús, M.I., Delgado, L.C., Perakakis, P., Kley, E., Mata, J.L., and Rodríguez, S. (2007). Cardiac defense: From attention to action. Int. J. Psychophysiol. 66, 169–182.

36. O’Mara, M.T., Wikelski, M., Voigt, C.C., Maat, A. Ter, Pollock, H.S., Burness, G., Desantis, L.M., and Dechmann, D.K.N. (2017). Cyclic bouts of extreme bradycardia counteract the high metabolism of frugivorous bats. Elife 6, 1–20.

37. Graham, F.K., and Clifton, R.K. (1966). Heart-rate change as a component of the orienting response. Psychol. Bull. 65, 305–320.

38. Shimizu, H., and Okabe, M. (2007). Evolutionary origin of autonomic regulation of physiological activities in vertebrate phyla. J. Comp. Physiol. A Neuroethol. Sensory, Neural, Behav. Physiol. 193, 1013–1019.

39. Shuranova, Z. evidencia SN autonoma en decapodos.pdf.

40. Dulcis, D., and Levine, R.B. (2003). Innervation of the heart of the adult fruit fly, Drosophila melanogaster. J. Comp. Neurol. 465, 560–578.

41. Dulcis, D., Levine, R.B., and Ewer, J. (2005). Role of the neuropeptide CCAP in Drosophila cardiac function. J. Neurobiol. 64, 259–274.

42. Rickenbacher, E., Perry, R.E., Sullivan, R.M., and Moita, M.A. (2017). Freezing suppression by oxytocin in central amygdala allows alternate defensive behaviours and mother-pup interactions. Elife 6, 1–17.

43. Verma, D., Wood, J., Lach, G., Herzog, H., Sperk, G., and Tasan, R. (2016). Hunger promotes fear extinction by activation of an amygdala microcircuit. Neuropsychopharmacology 41, 431–439.

44. Vale, R., Evans, D.A., and Branco, T. (2017). Rapid Spatial Learning Controls Instinctive Defensive Behavior in Mice. Curr. Biol. 27, 1342–1349. Available at: http://dx.doi.org/10.1016/j.cub.2017.03.031.

45. Eilam, D. (2005). Die hard: A blend of freezing and fleeing as a dynamic defense - Implications for the control of defensive behavior. Neurosci. Biobehav. Rev. 29, 1181–1191.

46. Lebestky, T., Chang, J. S., Dankert, H., Zelnik, L., Kim, Y. C., Han, K. A., Wolf, F. W., Perona, P., and Anderson, D. J. Two different forms of arousal in Drosophila are oppositely regulated by the dopamine D1 receptor ortholog DopR via distinct neural circuits. Neuron, 64(4), 522–536 (2009).

47. Appelhans, B.M., and Luecken, L.J. (2006). Heart rate variability as an index of regulated emotional responding. Rev. Gen. Psychol. 10, 229–240.

48. Sacha, J. (2014). Interaction between heart rate and heart rate variability. Ann. Noninvasive Electrocardiol. 19, 207–216.

49. López, R., Poy, R., Pastor, M.C., Segarra, P., and Moltó, J. (2009). Cardiac defense response as a predictor of fear learning. Int. J. Psychophysiol. 74, 229–235. Available at: http://dx.doi.org/10.1016/j.ijpsycho.2009.09.006.

50. Critchley, H.D., and Garfinkel, S.N. (2015). Interactions between visceral afferent signaling and stimulus processing. Front. Neurosci. 9, 1–9.

51. Azevedo, R.T., Badoud, D., and Tsakiris, M. (2018). Afferent cardiac signals modulate attentional engagement to low spatial frequency fearful faces. Cortex 104, 232–240.

52. Lo, P.C.H., and Frasch, M. (2001). A role for the COUP-TF-related gene seven-up in the diversification of cardioblast identities in the dorsal vessel of Drosophila. Mech. Dev. 104, 49–60.

53. Peirce, J.W. (2007). PsychoPy-Psychophysics software in Python. J. Neurosci. Methods 162, 8–13. Available at: http://dx.doi.org/10.1016/j.jneumeth.2006.11.017.

