## Supplementary data for "Threat induces changes in cardiac activity and metabolism negatively impacting survival in flies"

**Supplementary Materials**

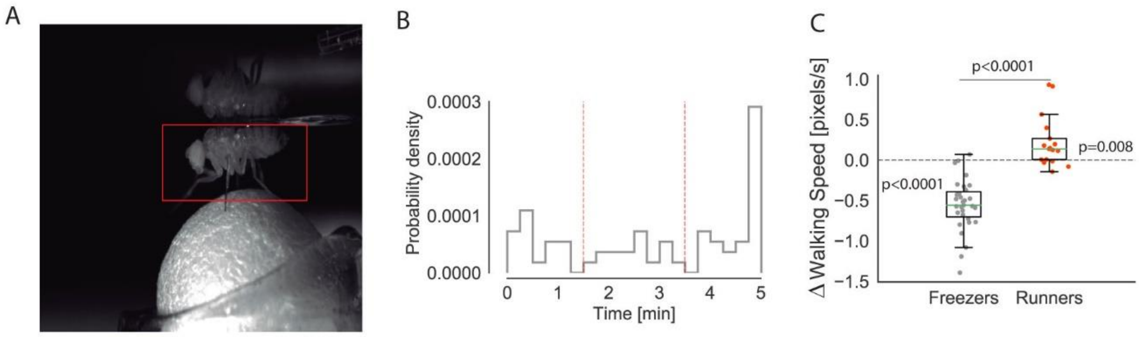


**Figure S1.** (A) Still image showing the ROI surrounding the fly used for the analysis of freezing. (B) Probability density distribution of time spent freezing by individual flies. Vertical red lines indicate threshold for runner flies, to the left of the first line, and freezer flies, to the right of the second line. (C) Change in walking speed per individual flies (each data point shows average speed during stimulation – average speed during baseline period). Green horizontal lines represent medians, box represents interquartile range and whiskers min and max values.

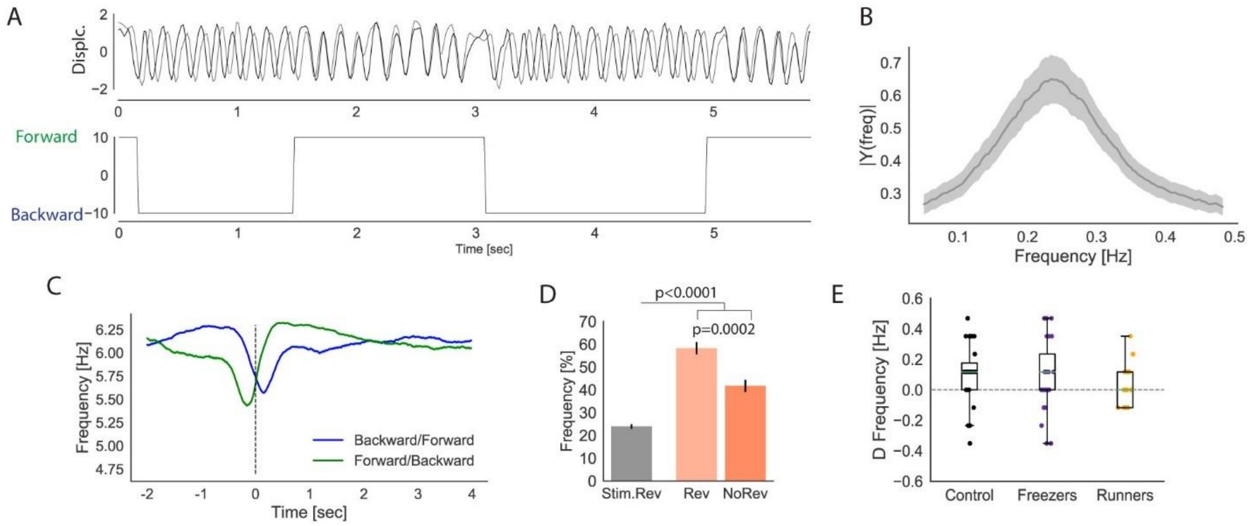


**Figure S2. (A)** Representative traces of the beating of anterior (black) and posterior (grey) cardiomyocyte nuclei (above). Square wave showing cardiac reversal. Forward beating is represented with the maximum value and backward with the minimum. **(B)** Average power spectral density of baseline cardiac reversal frequency (± s.e.m.) for stimulated flies. **(C)** Average peak beating frequency (± s.e.m.), aligned on reversal events. Dashed vertical lines represents directional change. **(D)** Frequency of cardiac arrest associated with looming with and without cardiac reversal (± s.e.m.). In grey, frequency of cardiac arrest associated with cardiac reversal in random points during stimulation period excluding points of looming presentation. **(E)** Change in the peak rate of forward beating per individual fly (stimulation – baseline period).

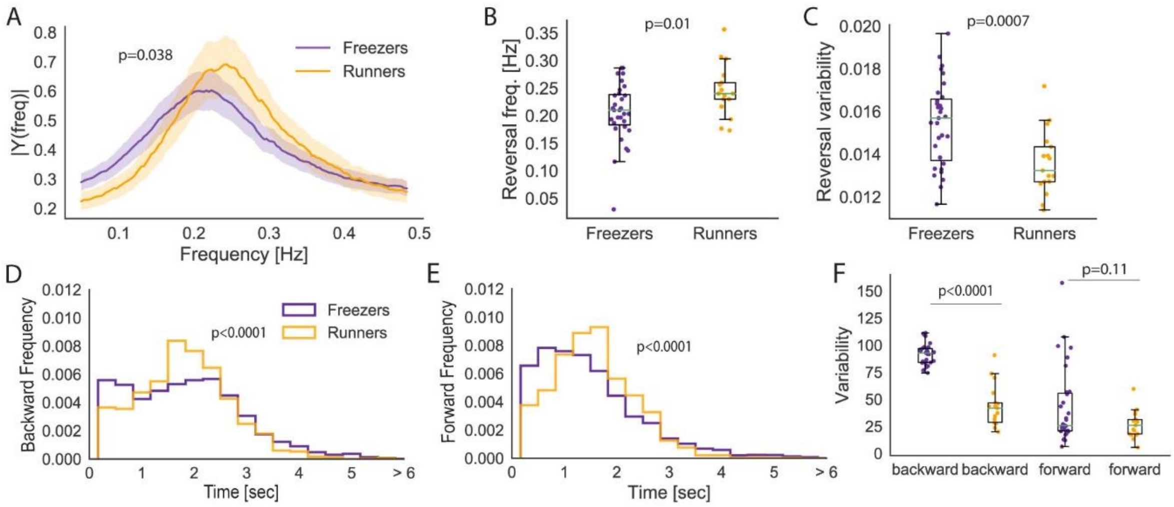


**Figure S3. (A)** Average power spectral density of cardiac reversal frequency during baseline period for freezers and runners. **(B)** Peak cardiac reversal frequency per individual fly during baseline period. **(C)** Cardiac reversal variability per individual fly during baseline period. **(D-E)** Density distribution of the length of backward (D) and forward bouts (E) during baseline period. In A-E, purple and orange represent freezer and runner flies. In B and C, green horizontal lines represent medians, box represents interquartile range and whiskers min and max values.

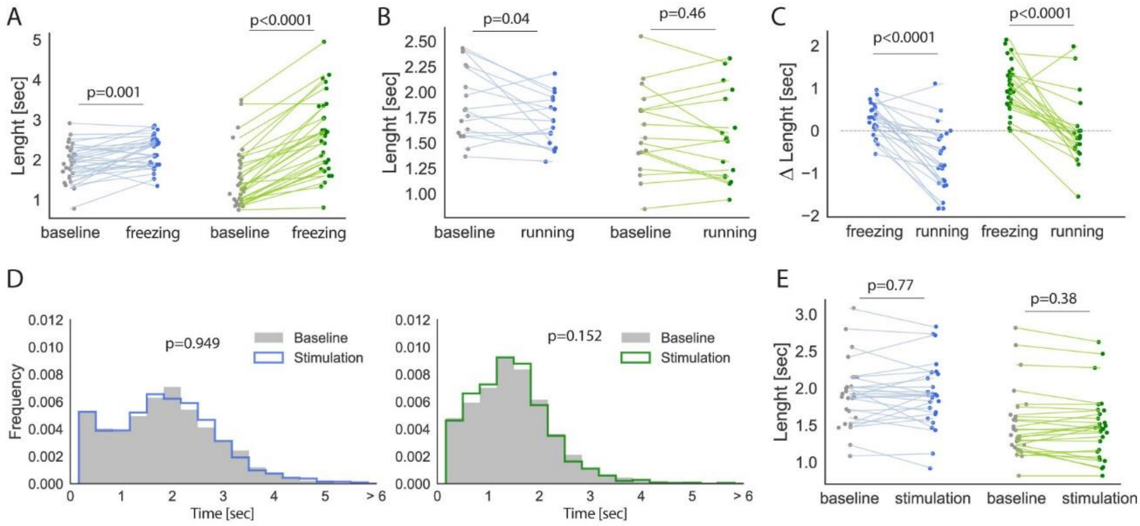


**Figure S4.** (A) Median length of backward and forward bouts per individual freezer fly during baseline and freezing periods. **(B)** Median length of backward and forward bouts per individual runner fly during baseline and running periods. **(C)** Change in the median length of backward and forward bouts per individual freezer fly (each data point shows median length during stimulation – median length during baseline period). **(D)** Density distribution of backward (left) and forward bouts length (right) during baseline and stimulation periods for control stimulated flies. **(E)** Median length of backward and forward bouts per individual control fly. In A-E, blue and green correspond to backward and forward bouts respectively.

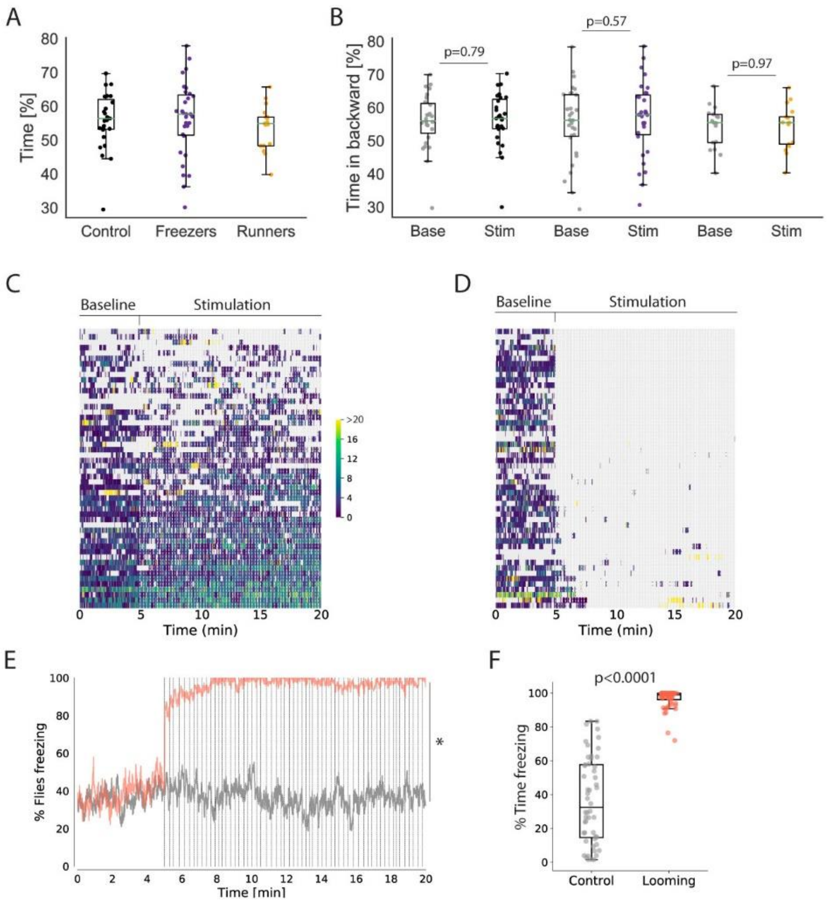


**Figure S5. (A)** Percentage of time in backward mode per individual fly during baseline period. **(B)** Percentage of time in backward mode per period of walking during baseline (grey) or during stimulation in control (black), freezer (purple) and runner (orange) flies. **(C-D)** Walking speed and freezing/immobility (grey) in 1 sec bins for control (C, n=52) and looming stimulated flies (D, n=52). Each row corresponds to one fly, rank ordered by maximum average walking speed during stimulation period. **(E)** Fraction of flies freezing. **(F)** Percentage of time freezing per individual fly. In E and E, grey corresponds to control flies and red to looming stimulated flies. Dashed vertical lines in C, D, F and G represent stimulus presentations. E-F include walking bouts only. Green horizontal lines represent medians, box represents interquartile range and whiskers min and max values.

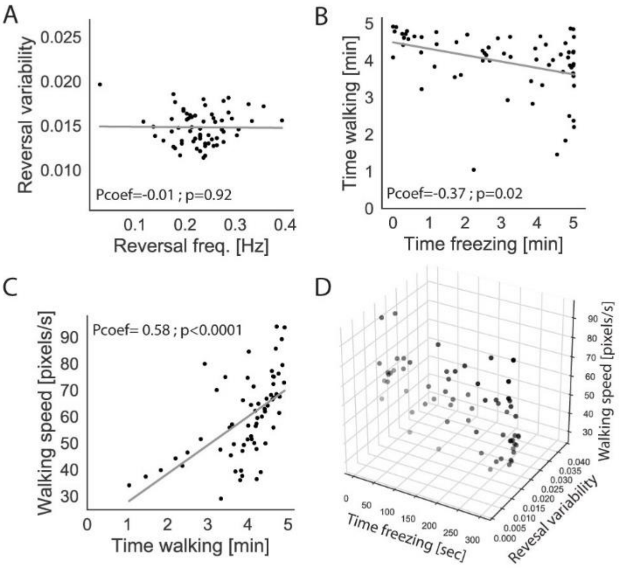


**Figure S6.** **(A)** Scatter plot showing the value of baseline peak reversal frequency and reversal variability for each loom stimulated fly. **(B-C)** Scatter plot showing the value of the total time walking during baseline period and the total time freezing (B) and baseline average walking speed (C) for each loom stimulated fly. **(D)** 3D scatter plot showing the value of total time freezing, baseline cardiac reversal variability and baseline walking speed for the 64 loom stimulated flies. Least squares regression lines are shown in grey. Pcoef shows Pearson coefficient.

**Supplementary video 1. Freezing response to looming of a semi-restrained fly.**

**Supplementary video 2. Fleeing response to looming of a semi-restrained fly.**

**Supplementary video 3. Heart activity of an intact fly.**
